# Pubertal Development Underlies Optimization of Inhibitory Control Through Specialization of Ventrolateral Prefrontal Cortex

**DOI:** 10.1101/2022.04.28.489931

**Authors:** Orma Ravindranath, Finnegan J. Calabro, William Foran, Beatriz Luna

## Abstract

Inhibitory control improves into young adulthood after specialization of relevant brain systems during adolescence. However, the biological mechanisms supporting this unique transition are not well understood. Given that adolescence is defined by pubertal maturation, we examined relative contributions of chronological age and pubertal maturation to inhibitory control development. 105 8-19-year-olds completed 1-5 longitudinal visits (227 visits total) in which pubertal development was assessed via self-reported Tanner stage and inhibitory control was assessed with an in-scanner antisaccade task. As expected, percent and latency of correct antisaccade responses improved with age and pubertal stage. When controlling for pubertal stage, chronological age was distinctly associated with correct response rate. In contrast, pubertal stage was uniquely associated with antisaccade latency even when controlling for age. Chronological age was associated with fMRI task activation in several regions including the right dorsolateral prefrontal cortex, while puberty was associated with right ventrolateral prefrontal cortex (VLPFC) activation. Furthermore, task-related connectivity between VLPFC and cingulate was associated with both pubertal stage and response latency. These results suggest that while age-related developmental processes may support maturation of brain systems underlying the ability to inhibit a response, puberty may play a larger role in the effectiveness of generating cognitive control responses.

## Introduction

Cognitive control continues to improve through adolescence in parallel with the maturation of relevant brain systems, leading to its stability in adulthood (Luna et al., 2015). Specifically, while the ability to exert inhibitory control exists as early as infancy (Johnson, 1995), the rate of correct inhibitory responses significantly increases through adolescence, followed by a shift towards stabilization and optimization in adulthood (Bari and Robbins, 2013; Dempster, 1992; Luna, 2009; Luna et al., 2004; Ordaz et al., 2013). These developmental improvements are not linear, but rather, show rapid growth through childhood followed by a deceleration through adolescence and reaching a plateau in young adulthood (Luna et al., 2004; Ordaz et al., 2013). Similarly, prior work examining age-related changes in brain function during inhibitory control has shown that the dorsolateral prefrontal cortex (DLPFC) is predominantly engaged in childhood, but by adolescence, DLPFC activation decreases to adult levels, with engagement of the anterior cingulate cortex (ACC) mediating improvements in performance (Ordaz et al., 2013) as top-down frontal connectivity becomes established (Hwang et al., 2016, 2010). Thus, both changes in behavior and brain function show that developmental processes occurring uniquely in adolescence underlie the transition to adult-level effective and stable engagement of widely distributed executive systems supporting cognitive control.

Previous studies have examined cognitive development predominantly as a function of chronological age, limiting the ability to assess biological processes unique to the adolescent period and examine variability in developmental trajectories. Characterizing the mechanistic and neurobiological contributions to these neurocognitive processes is critical to understanding the establishment of adult trajectories, including the emergence of psychiatric disorders (e.g., psychosis, mood disorders, substance use disorders) during adolescence and young adulthood (Chambers et al., 2003; Paus et al., 2008; Pfeifer et al., 2011), many of which are associated with cognitive dysfunction (Castellanos-Ryan et al., 2014; Frost et al., 1989; Toren et al., 2000).

The beginning of adolescence is defined by the onset of puberty, when major changes in hormonal levels trigger developmental processes which underlie the transition into adulthood and are believed to play a role in critical period plasticity, which may be present in association cortex during adolescence (Larsen & Luna, 2018). Extensive research shows strong associations between pubertal hormones and cognitive function, underscoring the potential role of puberty in neurocognitive development (Almey et al., 2015; Bimonte and Denenberg, 1999; Colzato et al., 2010; Colzato and Hommel, 2014; Gibbs, 2005; Holmes et al., 2002; Ladouceur, 2012; Sandstrom et al., 2006; Vijayakumar et al., 2018). However, few studies have focused on puberty’s relationship with inhibitory control in particular. One recent study in a sample of 12-14-year-old healthy adolescents found no association between Tanner stage or testosterone levels and inhibitory control error rates, but did find that more advanced Tanner stage was associated with faster response latencies among females (Ordaz et al., 2018). Another study using drift-diffusion modeling found that the interaction of puberty and sex was significantly related to the amount of information considered to make decisions during an inhibitory control task (Castagna and Crowley, 2021). Less, if any, research exists on the associations between pubertal development and changes in brain activation and task-based connectivity during inhibitory control. However, disentangling pubertal effects from chronological age is challenging given their significant collinearity. Previous studies have separated these effects using statistical methods such as Akaike’s Information Criterion (AIC), (Akaike, 1974) to determine which measure produces the best-fitting model when associated with structural brain development or changes in resting state functional connectivity (Goddings et al., 2014; van Duijvenvoorde et al., 2019). Others have included age as a covariate when modeling the effect of puberty on structural brain development to examine the influence of pubertal development independent of age (Herting et al., 2017; Vijayakumar et al., 2021). Importantly, these studies incorporated longitudinal data, which is important for modeling both age- and puberty-related trajectories, and spanned wide age ranges incorporating the full span of adolescence (beginning at 7-8 years old and ending at 18-20 years old), so that all pubertal stages could be captured in a substantial number of individuals. Thus, in this study, we examined behavioral and fMRI data during an inhibitory control task using an accelerated longitudinal cohort of 8-19-year-olds, applying similar statistical methods (AIC, effect size, age covariate) to distinguish the effects of puberty and age on inhibitory control development and its underlying neurobiological correlates.

## Methods

### Participants

Data were acquired as part of a large, accelerated longitudinal study, in which neuroimaging and behavioral data were collected on 160 participants (8-30 years old) across 571 visits (1-5 visits per participant). Participants were recruited from the community and were screened for psychiatric and neurological problems in themselves and their first-degree relatives, as well as MRI contraindications such as metal in the body. For all subjects under the age of 18, parental consent and assent from the participant were obtained before beginning data collection. For those over the age of 18, consent was obtained from the participant. Over the course of the study, 24 individuals (14 females) did not complete follow-up visits due to obtaining braces, issues with scheduling or contacting, loss of interest, and change of residence. All experimental procedures were approved by the University of Pittsburgh Institutional Review Board and complied with the Code of Ethics of the World Medical Association (Declaration of Helsinki, 1964).

Because this study examines the effects of pubertal development, participants without pubertal data were excluded from all analyses. In order to measure only ongoing pubertal development, we also excluded participants who were already in Stage 5 (completed puberty) at their first study visit. The final sample used in all analyses included behavioral data for 227 study visits (1-5 repeated visits) from 105 participants (53 female, 8.0-19.3 years old, Figure 1) and fMRI data for 205 study visits from 98 participants (49 female) with additional exclusions due to participant sleepiness, excessive head motion, number of usable functional runs, lack of usable structural images, poor eye tracking quality, scanner inhomogeneities, and issues with data processing and recovery. We note that the subjects excluded due to data quality/issues did not differ significantly in age from the subjects included in the analyses.

**Figure 1:**
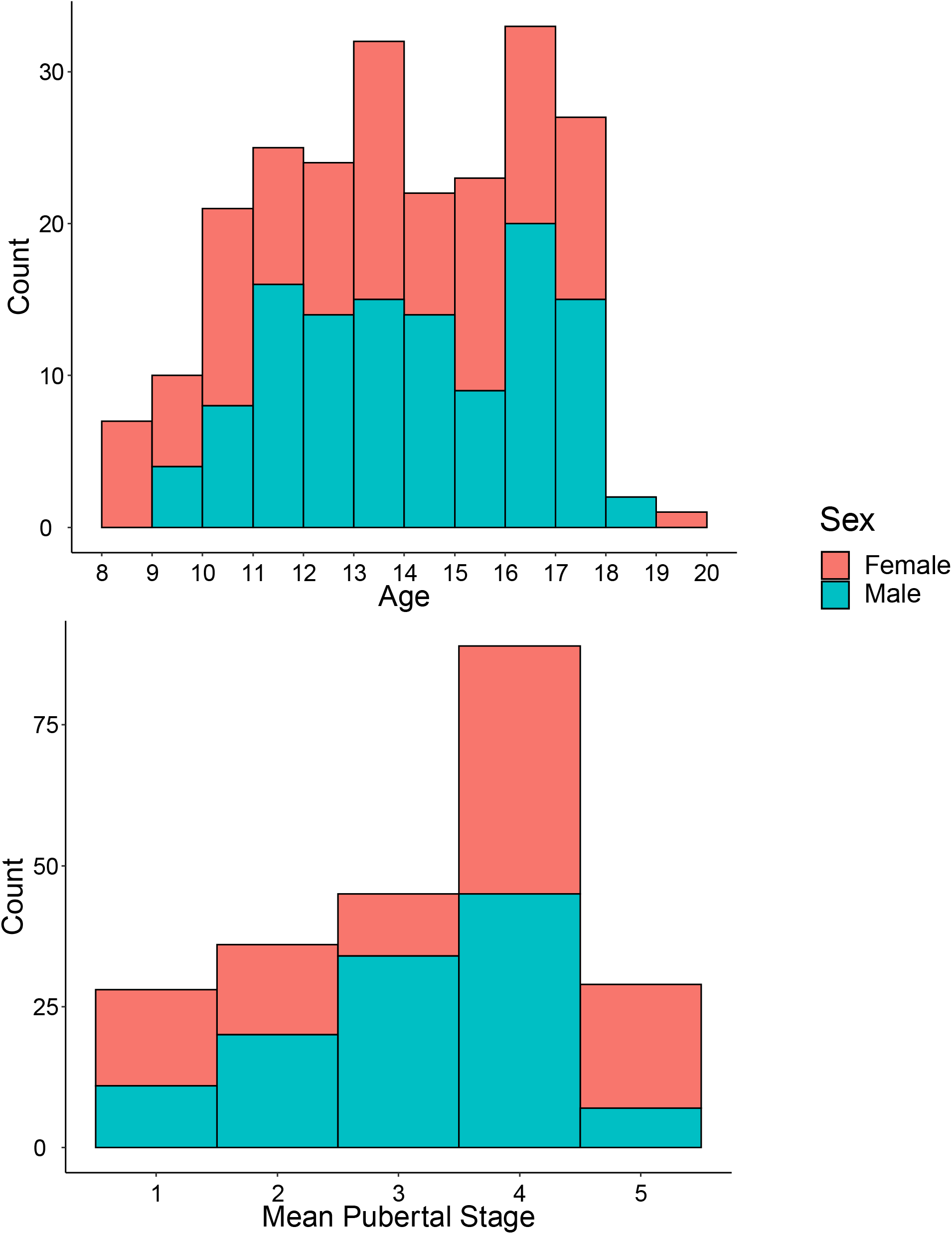
Distribution of included participants across age (top) and pubertal stage (bottom), colored by sex.

### Pubertal Assessments

Pubertal stage was assessed based on two self-report questionnaires. The first questionnaire was a pictorial Tanner staging questionnaire, in which line drawings of breast development, pubic hair growth, testes/scrotum/penis development, and testicular size were provided for Tanner Stages 1-5. Participants were asked to select the drawing that best corresponded with their own development at that time, which generated an overall Tanner score for the individual. The second questionnaire was the Pubertal Development Scale (Petersen et al., 1988), in which participants were asked questions about the development of various primary and secondary sexual characteristics, and chose from several options along the lines of “has not yet started growing”, “has barely started growing”, “is definitely underway”, and “seems completed”). The scoring of this questionnaire provides a pubertal stage on a 4-point scale, which was converted to a 5-point scale analogous to Tanner Stage using a previously-created evidence-based coding mechanism (Shirtcliff et al., 2009). For this study, the 5-point PDS score and the Tanner score were then averaged to create a Mean Pubertal Stage measure that was used to represent overall pubertal development in all analyses.

### Task Design

During fMRI scanning, participants completed an antisaccade (AS) task to assess inhibitory control (Hwang et al., 2010; Ordaz et al., 2013; Velanova et al., 2009, 2008). Full details of the experimental paradigm (Figure 2) are described by Velanova et al. (2008). Each run consisted of a block of the AS task and a block of the visually guided saccade (VGS) task (118.5 seconds each), preceded and separated by three blocked periods of fixation (36, 48, and 39 seconds each). Task order was counterbalanced across runs and participants. Twelve AS or VGS trials were presented in each task block, for a total of 48 of each trial type. Intertrial intervals (3–9 s) were pseudo-randomly distributed to permit estimation of trial-related activation (Dale, 1999). Each trial began as participants fixated on a colored cross-hair for 3 seconds instructing them to make a VGS (green) or an AS (red). Next, the saccade target stimulus, a yellow circle, appeared at one of six horizontal eccentricities (±3°, 6°, or 9°) for 1.5 s. For AS trials, participants were instructed to inhibit the reflexive saccade toward the target and to look instead to its horizontal mirror location. Target location order was randomized within each task block and no “gap” was interposed between the instruction cue and saccade target stimulus.

**Figure 2:**
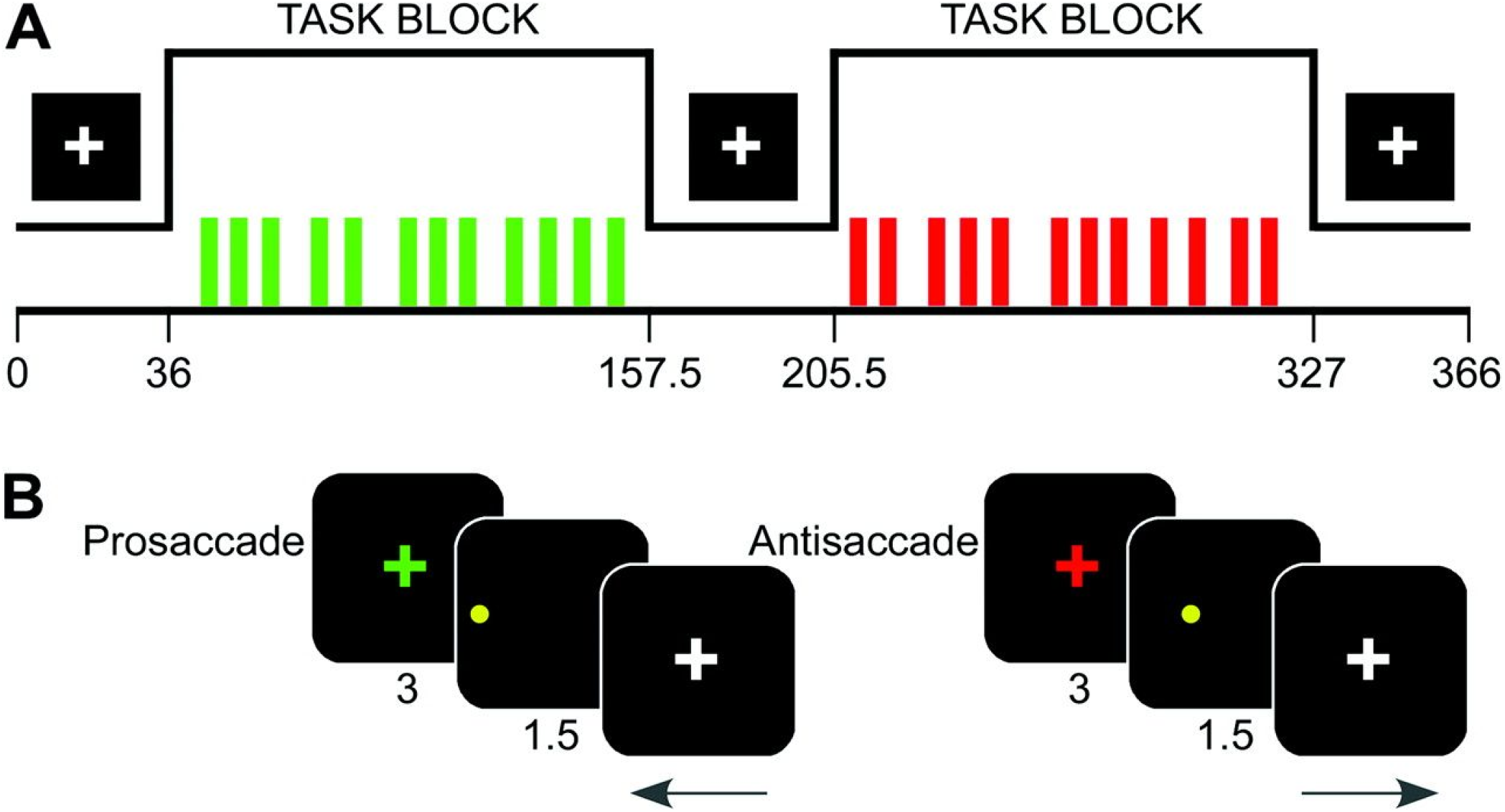
Diagram of experimental task paradigm, depicting structure of experimental run (A) and trial type (B). Originally from Velanova et al. (2008) and Ordaz et al. (2013).

### Eye Tracking Data Acquisition & Scoring

Eye movement measurements were obtained during fMRI scanning using a long-range optics eye-tracking system (Model R-LRO6, Applied Science Laboratories) with a sampling rate of 60 Hz. Nine-point calibrations were performed at the beginning of the session and between runs as necessary. Correct AS responses were defined as those in which the first eye movement during the saccade epoch with velocity greater than or equal to 30°/sec was made toward the mirror location of the peripheral cue and extended beyond a 2.5°/ visual angle from central fixation. Trials in which no eye movements were generated, gaze tracking was lost, blinks occurred before first onset, and express saccades (saccade starting within the first 4 samples (60Hz) after trial onset) occurred were excluded from analyses. This scoring system was automated using custom software which has been made publicly available on GitHub (https://github.com/LabNeuroCogDevel/autoeyescore). The two variables of interest used in subsequent analyses were the AS correct response rate (number of correct response trials / total number of usable trials) and mean response latency across correct trials.

### MR Data Acquisition

Data were acquired using a Siemens 3-Tesla MAGNETOM Allegra fitted with a standard circularity-polarized head coil. Head movement was minimized through use of pillows during scanning to immobilize the head and prior acclimation in an MR simulator. A PC (Dell Dimension 8200, Pentium 4, 2 GHz, Windows XP) running E-Prime (Psychology Software Tools) was used for displaying stimuli. Structural images were acquired using a sagittal magnetization-prepared rapid gradient-echo T1-weighted sequence (TR = 1570 ms; echo time [TE] = 3.04 ms; flip angle = 8°; inversion time [TI] = 800 ms; voxel size = 0.78125 × 0.78125 × 1 mm) and used for functional image alignment. Participants performed four functional runs (6 min 15 sec each), during which functional images were acquired using an echoplanar sequence sensitive to blood oxygen level-dependent contrast [T2*] with 29 contiguous 4-mm-thick axial images acquired parallel to the anterior–posterior commissure plane (TR = 1500 ms; TE = 25 ms; flip angle = 70°; voxel size = 3.125 × 3.125 mm in-plane resolution).

### fMRI Preprocessing & First-Level Analysis

Structural MRI data were preprocessed to extract the brain from the skull, and warped to the MNI standard brain using both linear (FLIRT) and non-linear (FNIRT) transformations. Preprocessing of task fMRI data followed our in-house standard protocol that incorporates tools from AFNI, NiPy, FSL and BrainWavelet (Cox, 1996; Gorgolewski et al., 2011; Jenkinson et al., 2012; Paulsen et al., 2015). The preprocessing pipeline included 4D slice-timing and head motion correction, skull stripping, intensity thresholding, wavelet despiking (Patel and Bullmore, 2016), co-registration to the structural image and nonlinear warping to MNI space, spatial smoothing using a 5-mm full width at half maximum Gaussian kernel, and intensity normalization. Source code for this pipeline is available online (https://github.com/LabNeuroCogDevel/fmri_processing_scripts).

First level analysis for task-related activation was performed by modeling all trial events in AFNI’s 3dDeconvolve (Cox, 1996). Events were defined by condition (visually-guided saccade or antisaccade) and participant’s performance on that trial (correct, corrected error, uncorrected error, or dropped) and modeled as a 4.5-second block. At this stage, nuisance regression was applied for the six head motion parameters, cerebrospinal fluid signal, and white matter signal. Volumes were censored if they contained framewise displacement (FD) estimates > 0.9 mm and subjects with greater than 15% of volumes censored were excluded from group-level analyses. To probe task-related brain activity, BOLD activation was then extracted from 13 inhibitory control regions-of-interest (ROIs), originally derived from Neurosynth (see Ordaz et al., 2013 for additional details of inhibitory control ROI selection), based on prior work showing that these regions are reliably engaged during inhibitory control. These regions encompass both those associated with the executive function component of inhibitory control (bilateral DLPFC, bilateral ventrolateral prefrontal cortex, dorsal ACC), and those associated with the motor control required for successful inhibitory control (supplementary eye field, pre-supplementary motor area, bilateral frontal eye field, bilateral parietal eye fields, and bilateral putamen).

### PPI Analysis

First level analysis for psychophysiological interactions (PPI) was performed by first generating a GAM-based hemodynamic response function. The BOLD time series for each seed region was then extracted, detrended, transposed, and deconvolved with the hemodynamic response function. An interaction regressor was then created by combining the deconvolved time series with a 1D timing file defining when task conditions occurred throughout the runs. These files were concatenated across runs, and task events were modeled in 3dDeconvolve, with the addition of the seed time series regressor and the interaction regressor.

Seeds for PPI connectivity analyses were selected from among those ROIs in which task-related activation was changing significantly with age or puberty. Only ROIs associated with executive function rather than those previously associated with motor control were considered for inclusion as PPI seeds. To ensure focused and hypothesis-driven analyses, we selected the seed with the greatest effect size (as measured by the conditional R-squared value) for each of age and puberty, among those for which age and puberty respectively were the best fitting model based on AIC/R^2^. Based on these criteria, the right DLPFC and ventrolateral prefrontal cortex (VLPFC) were selected, and these ROIs were defined based on the original set of ROIs derived from Ordaz et al., 2013.

### Statistical Analysis

Behavioral and task activation fMRI data were analyzed with linear mixed-effects models using the lme4 and lmerTest packages in R (Bates et al., 2014; R Core Team, 2020). Linear-mixed effects models were used to examine the fixed effects of age, puberty, and sex on error rate, latency, and fMRI beta values during the task. Subject was included in the model as a random effect. Outliers in correct response rate, response latency, and fMRI beta values greater than 3 standard deviations from the mean were excluded from respective analyses. Within this model, puberty, linear age, and inverse age were examined and the best fitting model was determined using both AIC and the conditional R-squared value (variance explained by both fixed and random effects combined). The main effects of age and puberty controlling for sex were also tested with Generalized Additive Mixed Models (GAMMs) and AIC values compared to confirm that age or puberty was the best-fitting model even when allowing for non-linearity.

Statistical analysis for the PPI analyses leveraged AFNI’s 3dLME to test for voxelwise effects of puberty and age separately, controlling for sex, based on whole brain connectivity maps generated for the selected PPI seed regions. This analysis was masked to only consider voxels with a 50% or greater probability of being gray matter in the MNI-152 template (Collins et al., 1999; Fonov et al., 2011, 2009). Significant clusters with main effects of puberty and age were identified and mean parameter estimates were subsequently extracted for these clusters using 3dROIStats, followed by *post hoc* testing to determine the direction of developmental effects and test for associations with antisaccade performance measures. Results were corrected for multiple comparisons using a combination of cluster size and voxel probability, with parameters determined through a Monte Carlo simulation using AFNI’s 3dFWHMx and 3dClustSim program on randomly generated data within the gray matter mask with the same smoothness as the group mean smoothness estimated from first-level residuals for each region. The autocorrelation function (ACF) option was used when running these scripts, which then specified the cluster size threshold applied with a single voxel threshold of *p*<0.05 (cluster size:169 voxels) or p<0.02 (cluster size: 70 voxels), for increased specificity, that was required to achieve a cluster-wise corrected *p*<0.05, based on current recommendations to prevent obtaining false positive clusters of connectivity (Chen et al., 2015).

## Results

### Development of Antisaccade Performance

Linear mixed models were used to examine the main and interaction effects of puberty, linear age, and inverse age, controlling for the effect of sex. We found significant effects of puberty (β = 0.38, p_FDR_ < 0.001), linear age (β = 0.48, p_FDR_ < 0.001), and inverse age (β = -0.46, p_FDR_ < 0.001) on antisaccade correct response rate, with AIC and R-squared values indicating that linear age produces the best fitting model for participants ages 8-19 (AIC_linearage_ = 528.42, AIC_inverseage_ = 534.48, AIC_puberty_ = 551.78, Figure 3A, Table 1). When examining models that include additive or interactive effects of age and puberty together, only age was significant across these models (p < 0.001) while puberty was not, indicating that the effects of increasing age drive change in correct response rate seen across this sample. We confirmed these results with GAMM analyses, which indicated a linear fit as the best model for the effect of age (p<0.001, Figure 3B), and the AIC values of the age and puberty GAMMs also indicate that age controlling for sex produced the best fitting model (AIC_age_ = -235.59, AIC_puberty_ = -208.16).

**Figure 3:**
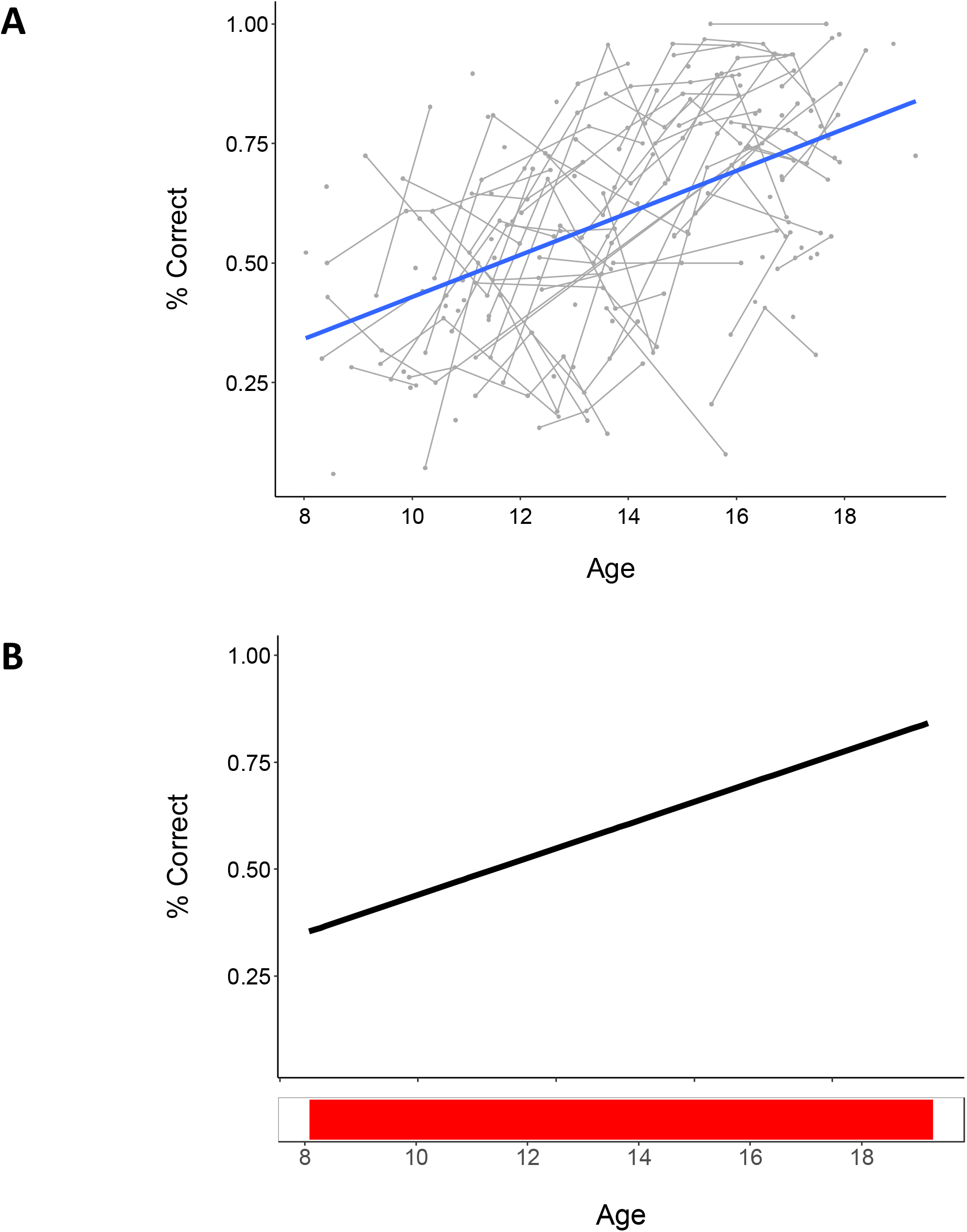
Plots of correct response rate data across age (years) using both a linear mixed model and generalized additive mixed model (B). Both show a significant positive association, such that increasing age is associated with improvements in correct response rate. Bottom panel in depicts intervals of significant age-related change. Individual datapoints reflect values for each session, and connected lines reflect sessions from the same participant.

**Table 1:** Regression statistics for linear mixed models assessing the main, additive, and interactive effects of puberty and age (linear and inverse forms) on correct response rate. All models are controlling for sex and statistics are FDR-corrected.

| Predictors | % Correct-Puberty |  |  | % Correct-Linear Age |  |  | % Correct-Inverse Age |  |  | % Correct-Puberty+Age |  |  | % Correct-Puberty*Age |  |  | % Correct-Puberty+Inverse Age |  |  | % Correct-Puberty*Inverse Age |  |  |
| --- | --- | --- | --- | --- | --- | --- | --- | --- | --- | --- | --- | --- | --- | --- | --- | --- | --- | --- | --- | --- | --- |
|  | Estimates | p | df | Estimates | p | df | Estimates | p | df | Estimates | p | df | Estimates | p | df | Estimates | p | df | Estimates | p | df |
| Intercept | -0.14 | 0.216 | 222.00 | -0.08 | 0.658 | 222.00 | -0.09 | 0.725 | 222.00 | -0.07 | 0.723 | 221.00 | -0.11 | 0.811 | 220.00 | -0.05 | 0.792 | 221.00 | -0.22 | 0.149 | 220.00 |
| Puberty | 0.38 *** | <0.001 | 222.00 |  |  |  |  |  |  | -0.08 | 0.723 | 221.00 | -0.03 | 0.811 | 220.00 | -0.03 | 0.792 | 221.00 | -0.00 | 0.978 | 220.00 |
| Sex | 0.25 | 0.170 | 222.00 | 0.07 | 0.658 | 222.00 | 0.06 | 0.725 | 222.00 | 0.06 | 0.771 | 221.00 | 0.07 | 0.811 | 220.00 | 0.04 | 0.792 | 221.00 | 0.09 | 0.750 | 220.00 |
| Age |  |  |  | 0.48 *** | <0.001 | 222.00 |  |  |  | 0.54 *** | <0.001 | 221.00 | 0.52 *** | <0.001 | 220.00 |  |  |  |  |  |  |
| Inverse Age |  |  |  |  |  |  | -0.48 *** | <0.001 | 222.00 |  |  |  |  |  |  | -0.48 *** | <0.001 | 221.00 | -0.57 *** | <0.001 | 220.00 |
| Puberty*Age |  |  |  |  |  |  |  |  |  |  |  |  | 0.03 | 0.811 | 220.00 |  |  |  |  |  |  |
| Puberty*Inverse Age |  |  |  |  |  |  |  |  |  |  |  |  |  |  |  |  |  |  | -0.15 * | 0.018 | 220.00 |
| Marginal R<sup>2</sup> / Conditional R<sup>2</sup> | 0.160 / 0.637 |  |  | 0.244 / 0.690 |  |  | 0.219 / 0.676 |  |  | 0.245 / 0.692 |  |  | 0.245 / 0.691 |  |  | 0.218 / 0.676 |  |  | 0.248 / 0.691 |  |  |
| AIC | 551.777 |  |  | 529.421 |  |  | 534.484 |  |  | 532.744 |  |  | 530.078 |  |  | 530.969 |  |  | 537.390 |  |  |

When testing the effects on response latency, we again found significant effects of puberty (β = -0.34, p_FDR_ < 0.001), linear age (β = -0.27, p_FDR_ < 0.001), and inverse age (β = 0.29, p_FDR_ < 0.001), but in this case, AIC and R-squared values indicated that puberty produced the best fitting model (AIC_linearage_ = 603.64, AIC_inverseage_ = 600.95, AIC_puberty_ = 594.33, Figure 4A, Table 2). Furthermore, the main effect of sex was also significant in the puberty model (β = -0.43, p_uncorrected_ =0.010, p_FDR_ = 0.030). For response latency, puberty and sex were significant throughout additive models of puberty and age (puberty controlling for the effect of age), while age was nonsignificant, indicating that unlike correct response rate, the development change in latency appears to be driven to a greater extent by pubertal maturation. We again confirmed these results with GAMM analyses, which indicated that pubertal development provided the best model fit (p<0.001, Figure 4B), and the AIC values of the age and puberty GAMMs also indicate that puberty produced the best fitting model (AIC_age_ = 2390.12, AIC_puberty_ = 2372.64). Notably, this GAMM model also indicated that the significant change in response latency occurred in pubertal stages 1-3.

**Figure 4:**
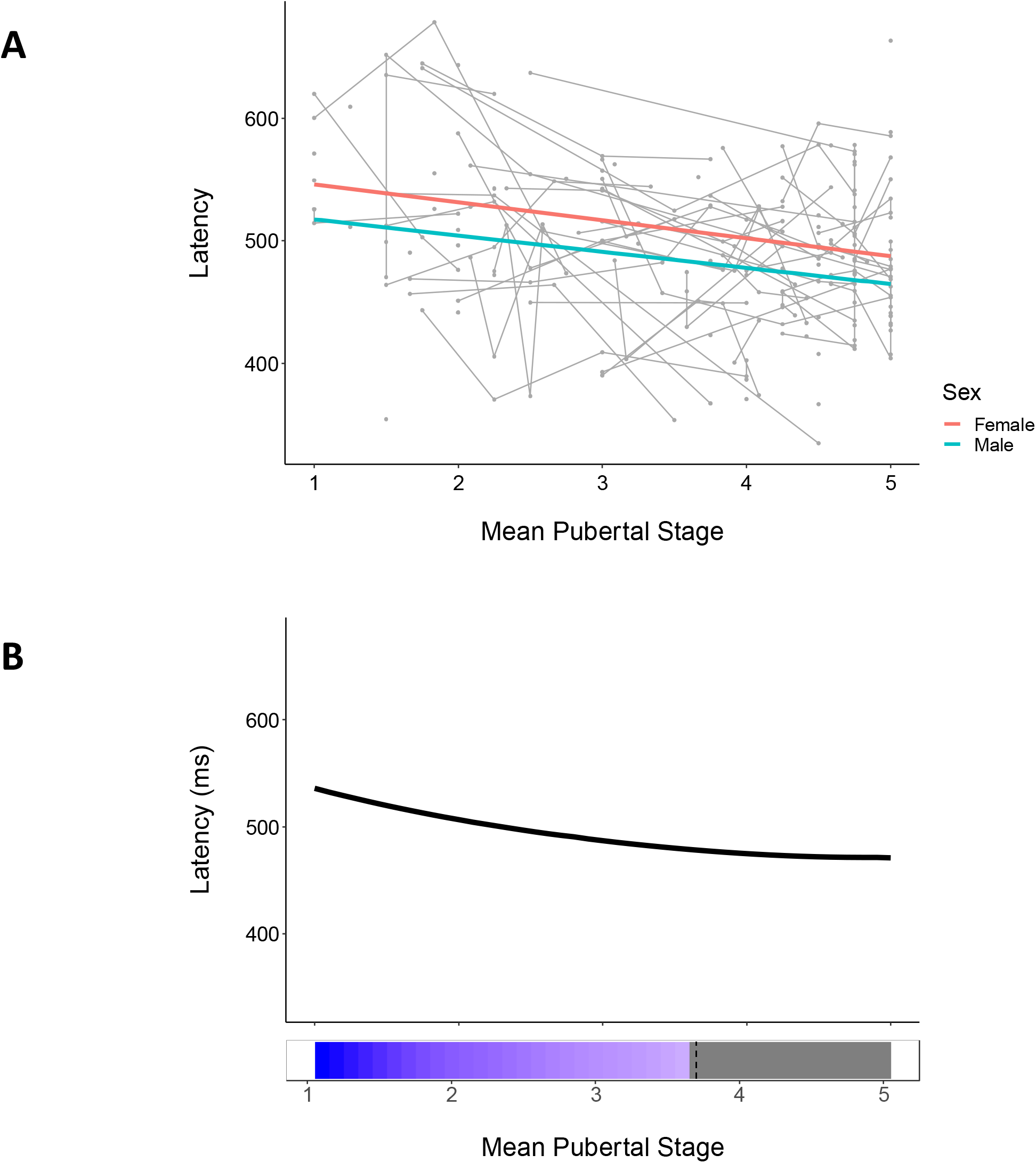
Plots of response latency data across mean pubertal stage using both a linear mixed model (A) and generalized additive mixed model (B). Both show a significant negative association, such that increasing pubertal stage is associated with decreasing response latency. Bottom panel in (B) depicts intervals of significant age-related change. Individual datapoints reflect values for each session, and connected lines reflect sessions from the same participant. In (A), red and blue lines reflect females and males, respectively.

**Table 2:**
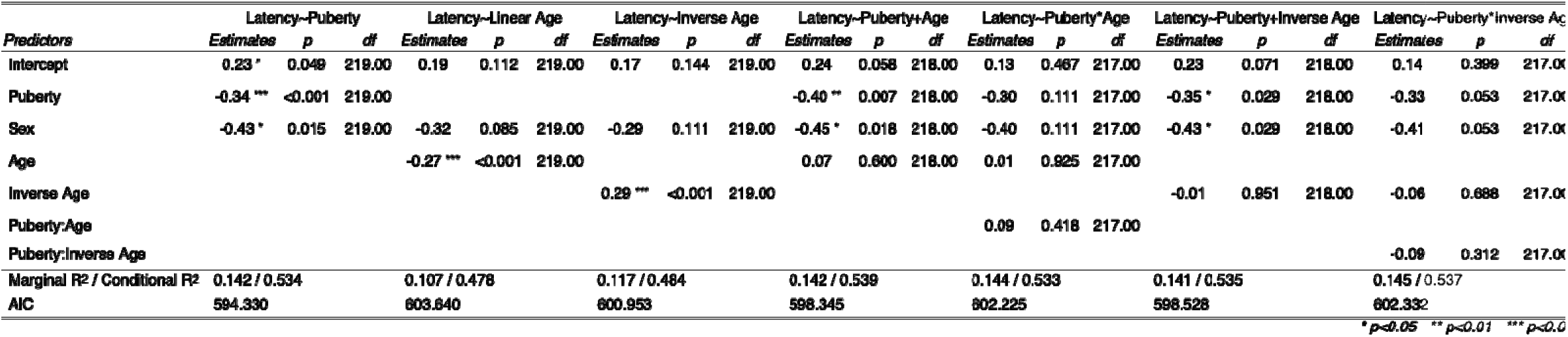
Regression statistics for linear mixed models assessing the main, additive, and interactive effects of puberty and age (linear and inverse forms) on response latency. All models are controlling for sex and statistics are FDR-corrected.

### Task Activation

Activation was measured within *a priori* regions of interest during performance of inhibitory control tasks (Ordaz et al., 2013). During correct trials, activation in the bilateral parietal eye fields, right DLPFC, right VLPFC, and dorsal ACC was significantly associated with both age and puberty (p_FDR_<0.05, Table 3, Figure 5) when tested separately. The inverse and/or linear age models were a better fit than puberty for all of these ROIs based on AIC and R^2^values, except for the right VLPFC, which was better fit by the puberty model (AIC_linearage_ = 569.67, AIC_inverseage_ = 569.55, AIC_puberty_ = 564.68). Activation in all ROIs that showed developmental effects was also tested for associations with correct response rate and latency, but all were nonsignificant at a threshold of p<0.05.

**Table 3:** Summary table of significant puberty, linear age, and inverse age associations with task activation extracted from 13 inhibitory control ROIs. Values in red are for the best-fitting model as per AIC and R^2^. L=left; R=right; SEF=supplementary eye field; pre-SMA=pre-supplementary motor area; FEF=frontal eye field; PEF=posterior eye fields; DLPFC=dorsolateral prefrontal cortex; VLPFC=ventrolateral prefrontal cortex; dACC=dorsal anterior cingulate cortex

|  | Puberty | Linear Age | Inverse Age |
| --- | --- | --- | --- |
| SEF | NS | NS | NS |
| Pre-SMA | NS | NS | NS |
| L FEF | NS | NS | NS |
| R FEF | NS | NS | NS |
| **L PEF** | **p=0.0322** | **p=0.0320** | **p=0.0326** |
| **R PEF** | **p=0.0123** | **p=0.00800** | **p=0.00458** |
| L DLPFC | NS | NS | NS |
| **R DLPFC** | **p=0.01312** | **p=0.00750** | **p=0.00458** |
| L VLPFC* | NA | NA | NA |
| **R VLPFC** | **p=0.00106** | **p=0.00750** | **p=0.00458** |
| **dACC** | **p=0.0383** | **p=0.0369** | **p=0.0216** |
| L Putamen | NS | NS | NS |
| R Putamen | NS | NS | NS |

**Figure 5:**
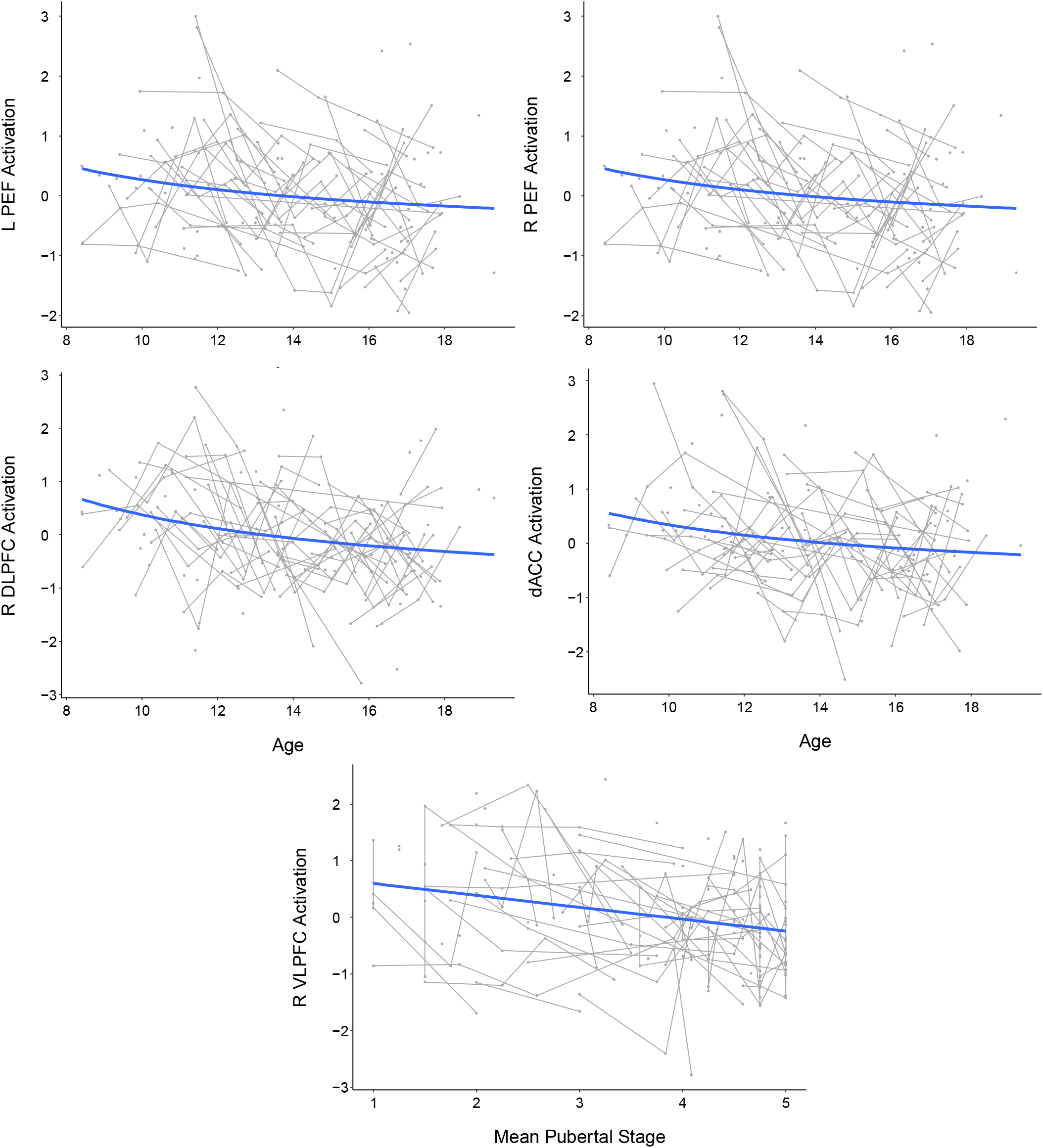
Plots of inhibitory control regions-of-interest (ROIs) in which significant age or puberty effects were observed. Both age and puberty were tested in each ROI, but the measure producing the best fitting model is visualized. Age was the most robust predictor of change in the bilateral parietal eye fields (PEF), right dorsolateral prefrontal cortex (DLPFC) and dorsal anterior cingulate cortex (dACC), while puberty was most associated with the right ventrolateral prefrontal cortex (VLPFC). Individual data points reflect values for each session, and connected lines reflect sessions from the same participant.

### Task-Related Connectivity

Because the association between pubertal development and task-related connectivity during inhibitory control has not been examined in the literature thus far, we conducted an exploratory PPI analysis based on the results of the behavioral and task activation analyses. Seed regions for the PPI analyses were selected from the developmentally- and task-relevant ROIs as described above. These analyses revealed, first, a widespread network of regions to which the right VLPFC was connected during correct antisaccade trials, including large areas across the frontal and parietal cortices, as well as regions of the striatum (Figure 6). Because we had identified puberty as the more robust predictor of task activation in the right VLPFC (as compared to age), we then tested for puberty-related developmental change across voxelwise task-related connectivity patterns seeded on the right VLPFC. This method identified significant clusters (FWE p<0.05) within the dACC, motor cortex, parahippocampal gyrus, and cingulate as areas for which task-related connectivity with the right VLPFC changed with pubertal development (Figure 7, Table 4). For each of these clusters, we tested for associations between mean connectivity strength across the cluster and response latency, since this was the behavioral measure more clearly associated with pubertal development. Notably, connectivity with the cingulate was significantly associated with latency (p_FDR_ = 0.0396), and remained significant when controlling for sex and puberty (Figure 8).

**Figure 6:**
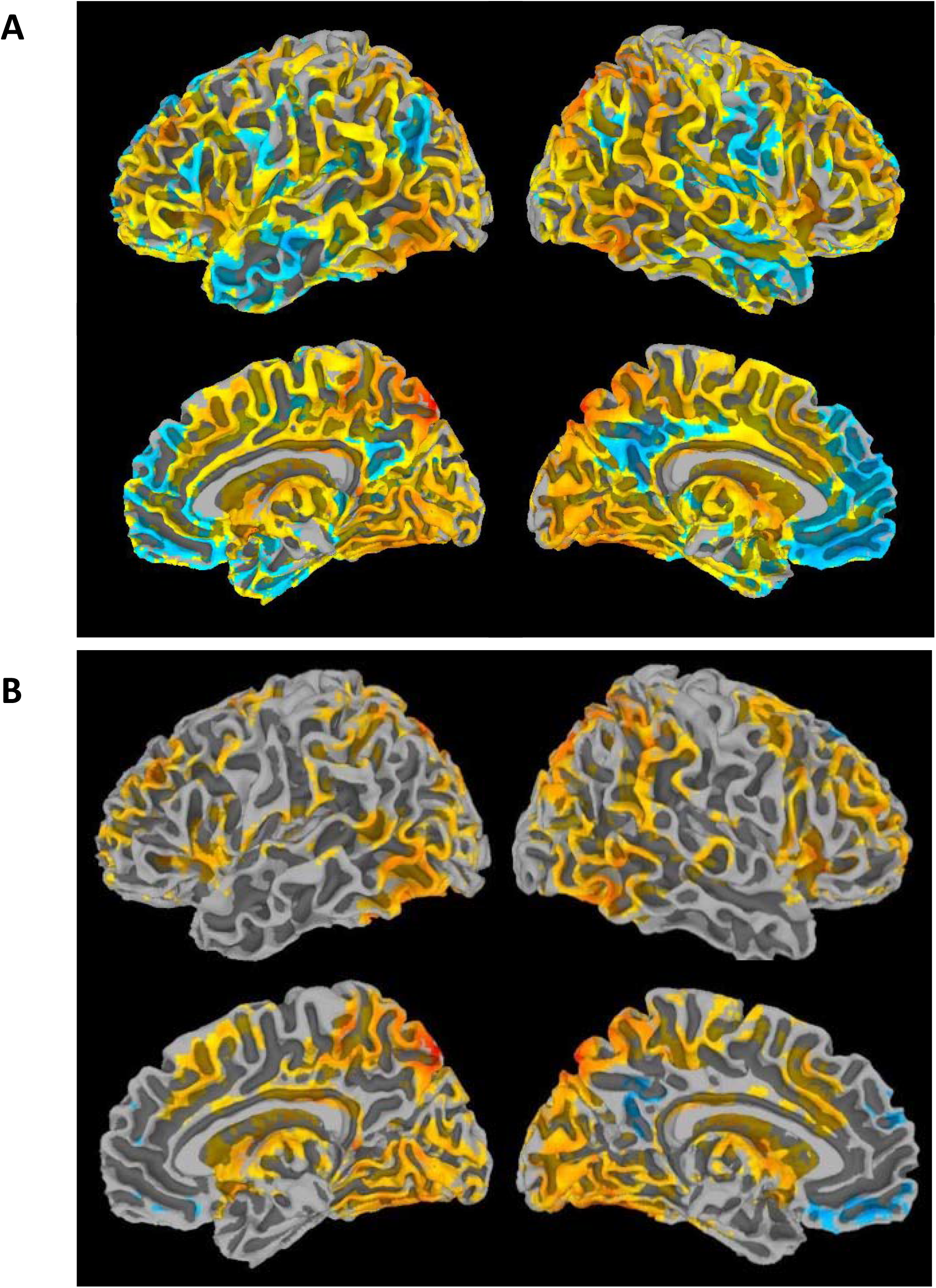
Maps showing R VLPFC PPI connectivity with the rest of the brain during correct trials. The top panel (A) shows unthresholded maps and the bottom (B) shows a voxelwise threshold of p_FDR_<0.05.

**Figure 7:**
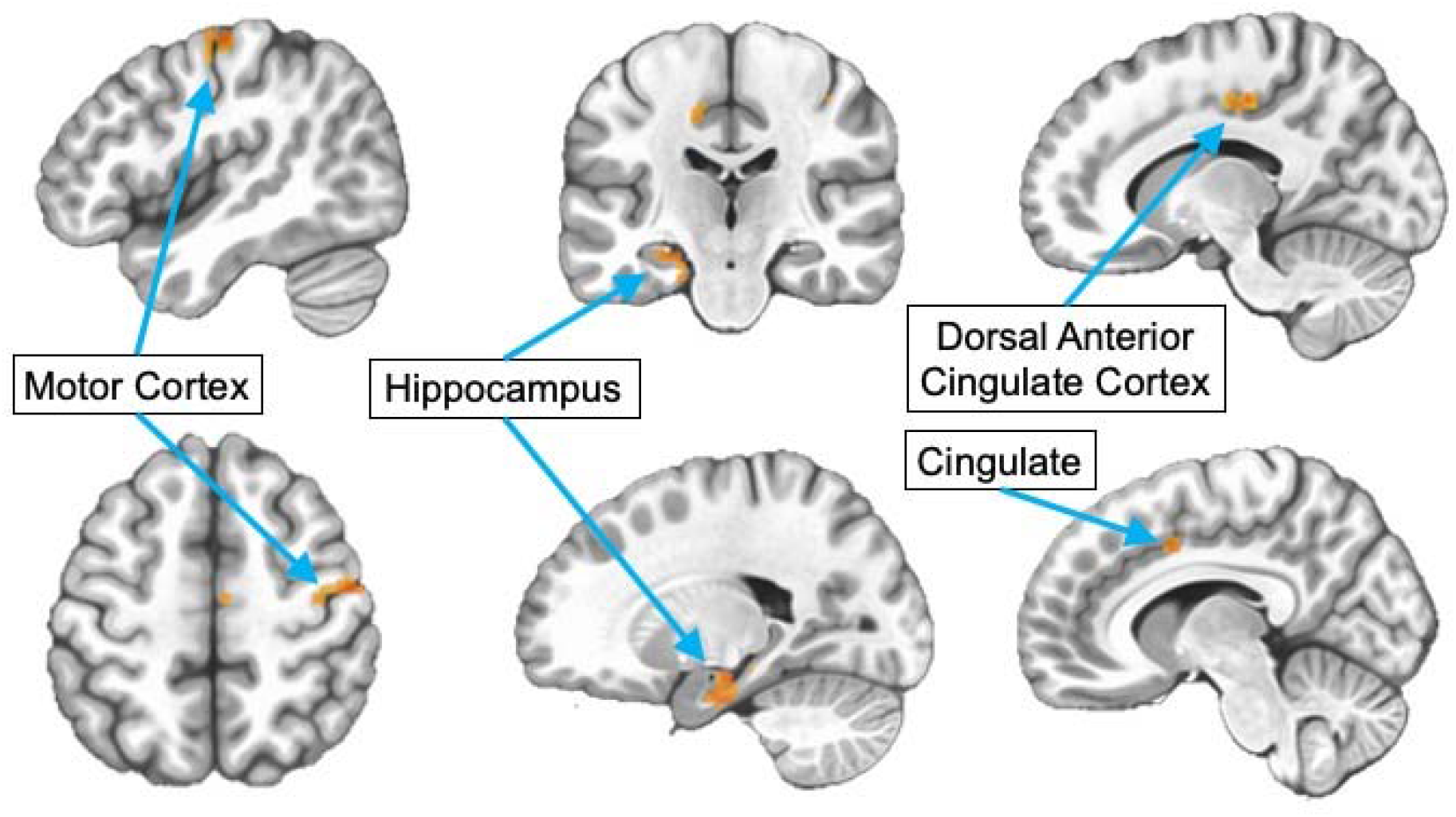
Maps showing clusters in R VLPFC PPI connectivity analysis that were significantly associated with pubertal stage when controlling for sex at a voxelwise threshold of p<0.02 (p<0.05 corrected).

**Figure 8:**
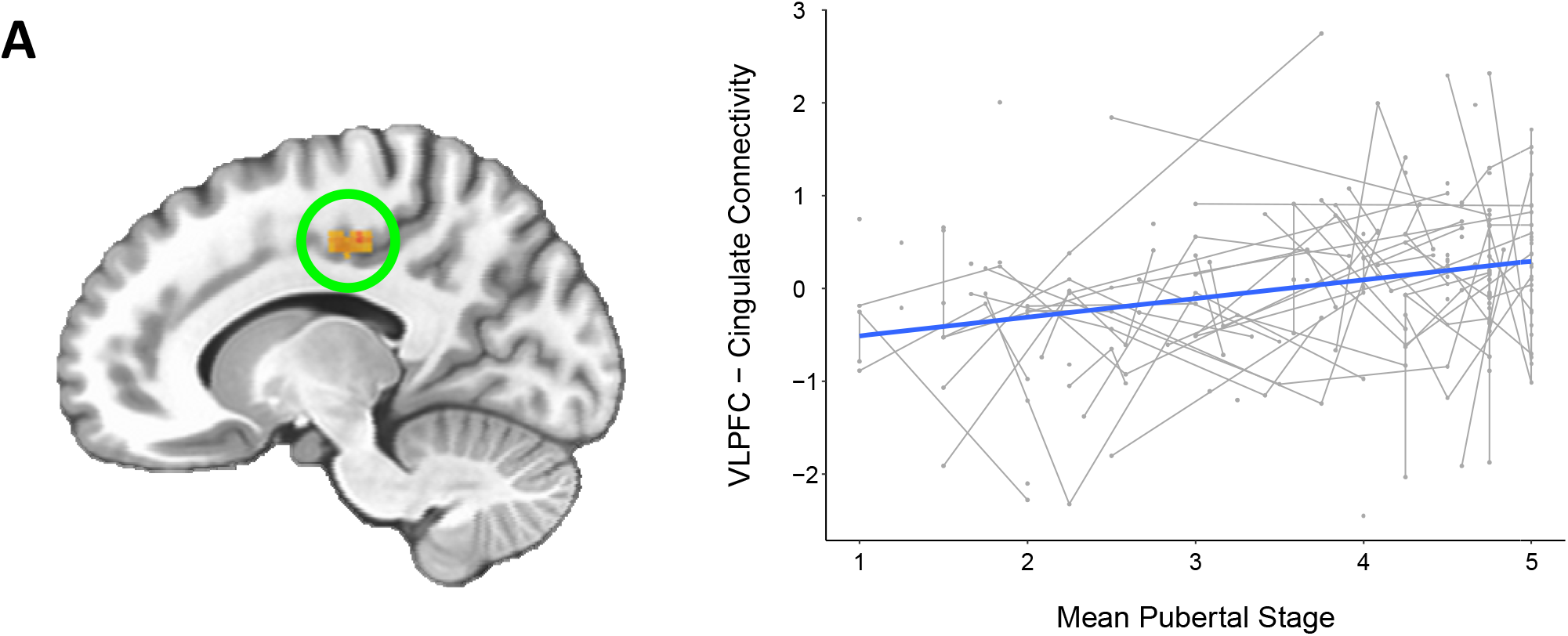
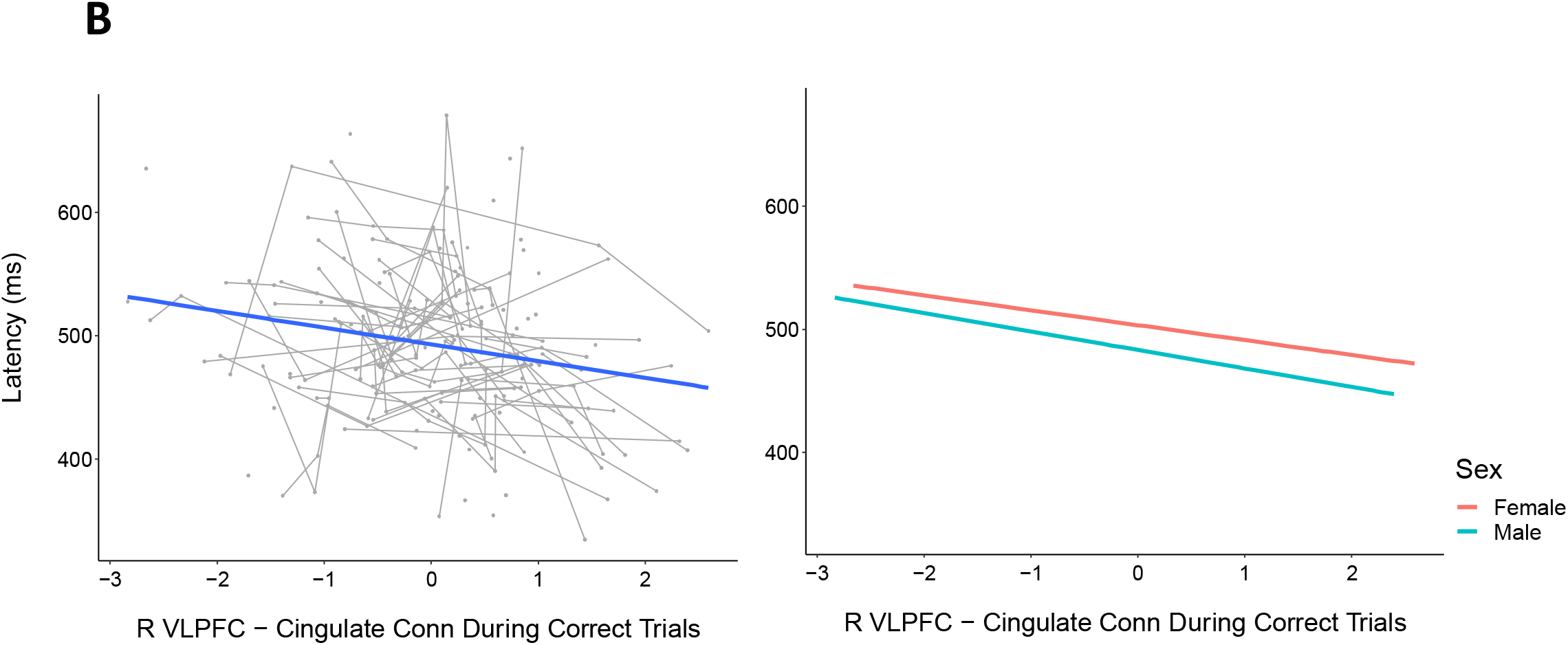
PPI analyses revealed several clusters within the brain in which task-related connectivity with the right VLPFC was significantly associated with pubertal stage, including the cingulate cluster visualized above (p<0.05 cluster corrected). Connectivity beta values were z-scored before plotting their association with mean pubertal stage (A). Within this cluster, connectivity was significantly associated with response latency, such that greater VLPFC connectivity was associated with faster response latencies (β = -0.19, p_FDR_ = 0.04). There was also a significant effect of sex such that females had slightly slower response latencies across this sample’s age range (β = -0.31, p = 0.03).

**Table 4:** Clusters for which right ventrolateral prefrontal cortex (VLPFC) connectivity is associated with pubertal stage at a voxelwise threshold of p<0.005 (p<0.05 cluster corrected). R = right; L = left; dACC = dorsal anterior cingulate cortex

| ROI # | Region | X | Y | Z | Cluster Size | Pos/Neg Effect |
| --- | --- | --- | --- | --- | --- | --- |
| 1 | R dACC | -5.8 | -8.4 | +39.8 | 95 vox | Positive |
| 2 | Motor Cortex | -49.4 | +7.7 | +53.6 | 94 vox | Positive |
| 3 | L Parahippocampal Gyrus | +17.2 | +16.9 | -17.6 | 91 vox | Positive |
| 4 | L Cingulate | +12.7 | +23.8 | +44.4 | 73 vox | Positive |

PPI analyses also revealed regions connected to the right DLPFC during correct trials, including areas of the prefrontal cortex, parietal cortex, and smaller portions of the striatum (Figure 9). Similar to the right VLPFC PPI analysis given above, we tested for connectivity changes with age since task activation in the right DLPFC was mostly strongly associated with chronological age, which revealed clusters in the cerebellum, motor cortex, superior temporal gyrus (STG) and inferior frontal gyrus (IFG) (Figure 10, Table 5). Among these regions whose DLPFC connectivity was changing with age, we examined associations with correct response rate, to determine if this connectivity might be a factor in the strong age-related component of this behavioral development. However, we did not find any significant associations between connectivity strength and antisaccade accuracy rates (all p_FDR_>0.05).

**Figure 9:**
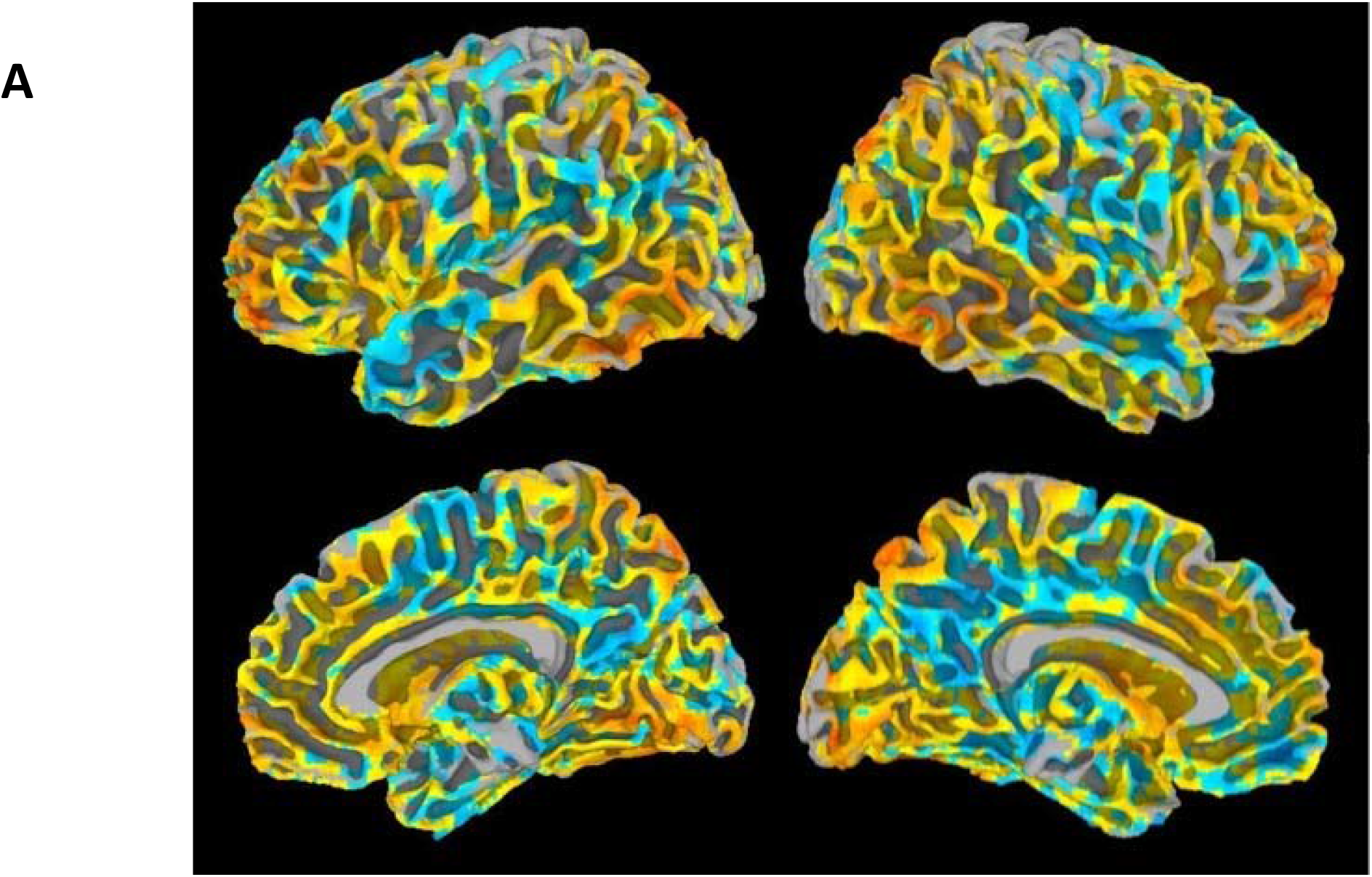
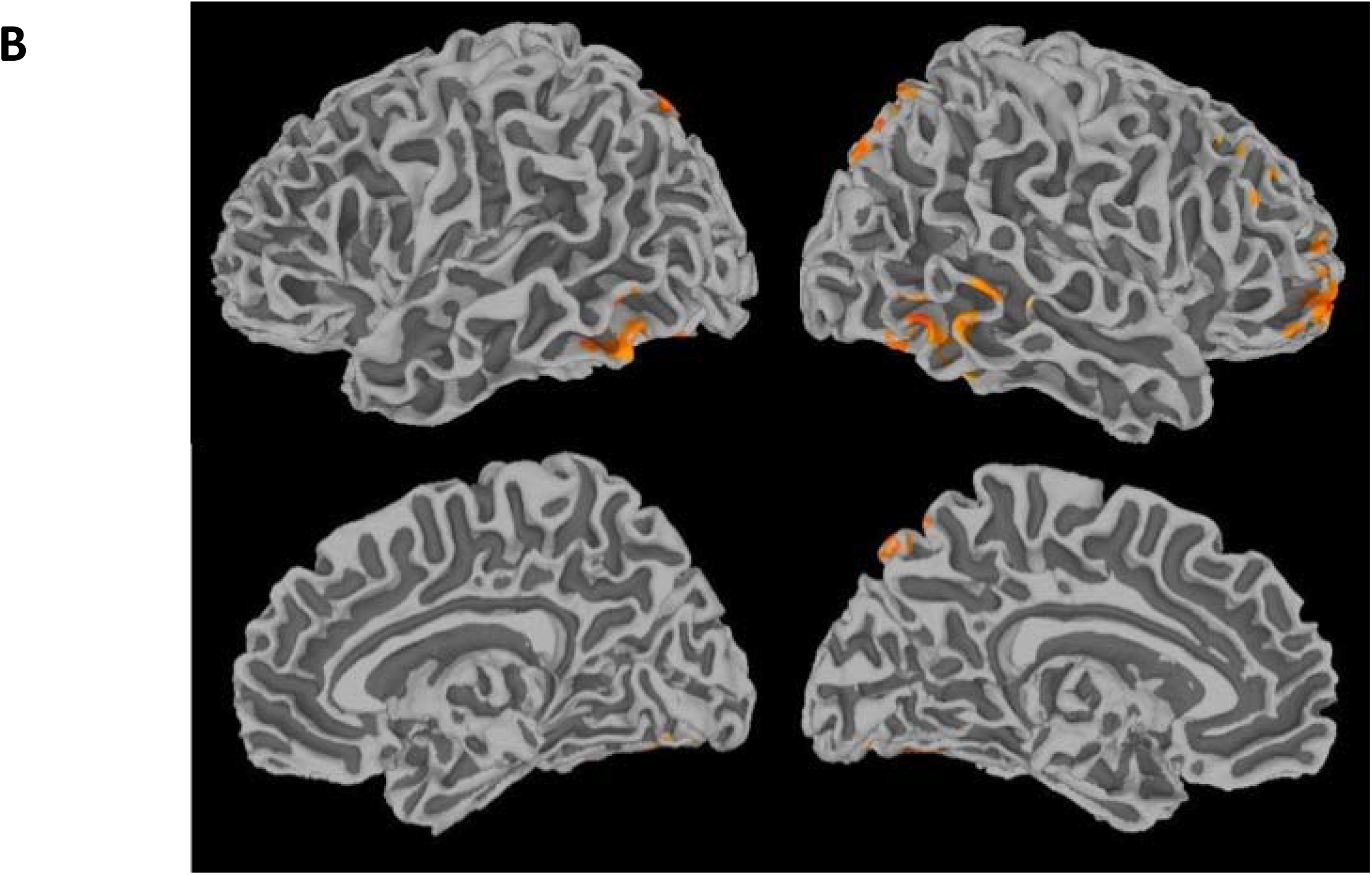
Maps showing R DLPFC PPI connectivity with the rest of the brain during correct trials. The top panel (A) shows unthresholded maps and the bottom (B) shows a voxelwise threshold of p_FDR_<0.05 (bottom)

**Figure 10:**
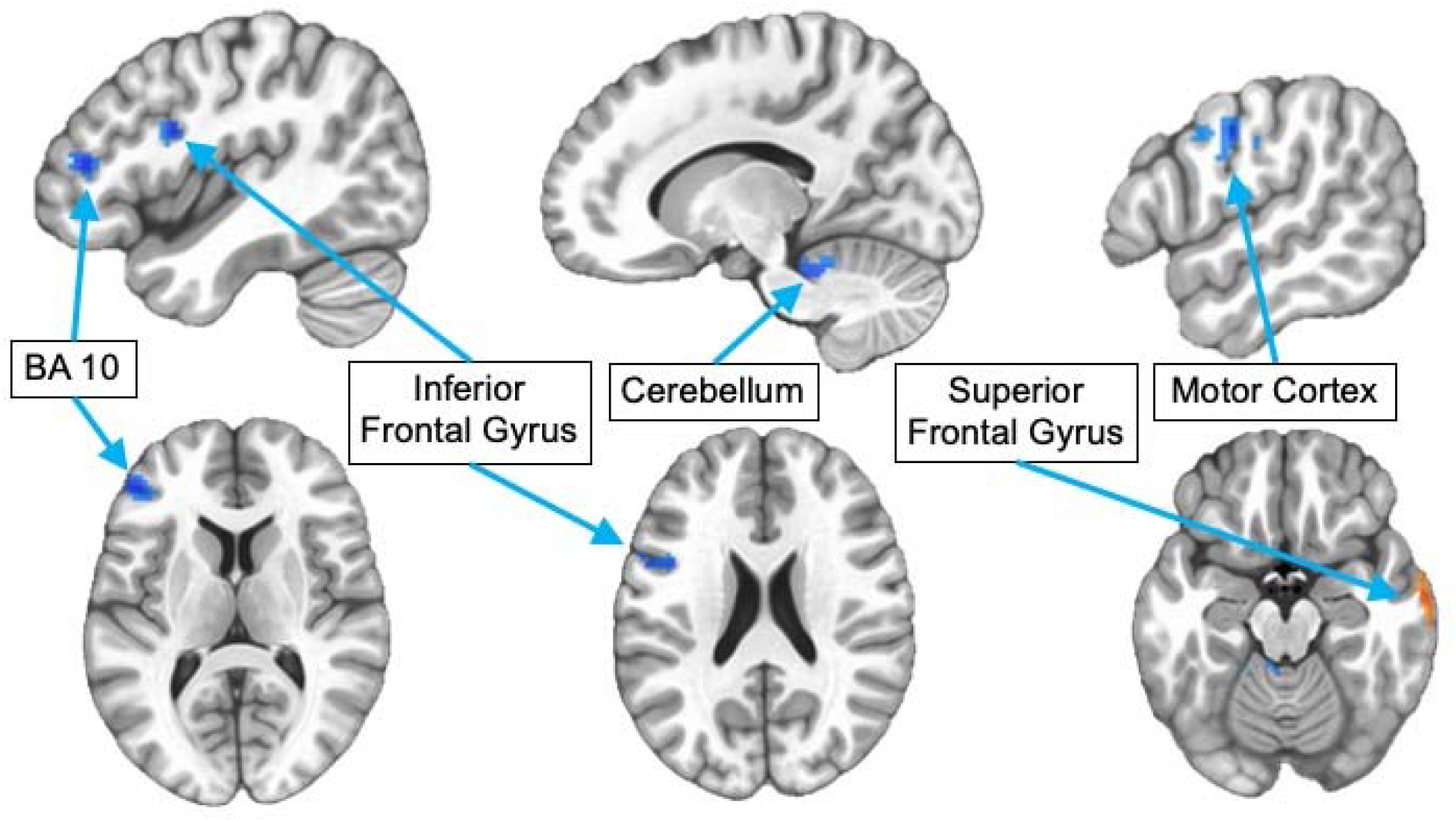
Maps showing clusters in R DLPFC PPI connectivity analysis that were significantly associated with chronological when controlling for sex at a voxelwise threshold of p<0.02 (p<0.05 corrected).

**Table 5:**
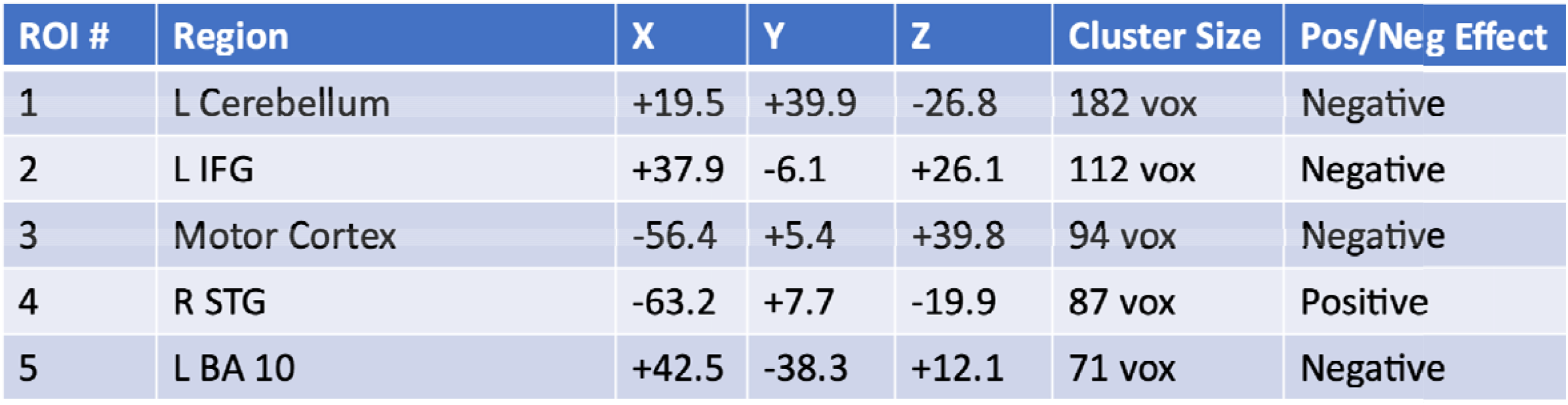
Clusters for which right dorsolateral prefrontal cortex (DLPFC) connectivity is associated with age at a voxelwise threshold of p<0.005 (p<0.05 cluster corrected). R = right; L = left; IFG=inferior frontal gyrus; STG=superior temporal gyrus; BA 10=Brodmann Area

| ROI # | Region | X | Y | Z | Cluster Size | Pos/Neg Effect |
| --- | --- | --- | --- | --- | --- | --- |
| 1 | L Cerebellum | +19.5 | +39.9 | -26.8 | 182 vox | Negative |
| 2 | L IFG | +37.9 | -6.1 | +26.1 | 112 vox | Negative |
| 3 | Motor Cortex | -56.4 | +5.4 | +39.8 | 94 vox | Negative |
| 4 | R STG | -63.2 | +7.7 | -19.9 | 87 vox | Positive |
| 5 | L BA 10 | +42.5 | -38.3 | +12.1 | 71 vox | Negative |

## Discussion

In this study, we examined unique associations between pubertal development and chronological age and inhibitory control, as well as associated neurobiological measures. We found that while development associated with chronological age drives improvements in correct response rate across adolescence, pubertal development was a more robust predictor of developmental improvement in response latency across this period. Furthermore, activity in the right VLPFC during correct inhibitory trials decreased with pubertal stage, and task-related connectivity of the right VLPFC to a region of the cingulate defined based on its association with pubertal stage was associated with developmental improvements in response latency. In contrast, the right DLPFC during correct inhibitory trials decreased with increasing age, with no particular task-related connectivity contributing to age-related change in inhibitory control performance.

A primary goal of this study was to understand the differential contributions of chronological age and puberty to the development of inhibitory control. However, an important consideration within this research question is what biological processes actually underlie development associated with chronological age, since age may encompass a large number of biological processes (including puberty). Indeed, some of this age-related change can simply be attributed to genetic and biological growth programming that drives fundamental human maturational processes. Another key driver that may interact with age-related processes is accumulated experience, which encompasses every day and stressful life events, the practice of skills, social interactions, and many other occurrences that shape how individuals perceive and process the world (Panchanathan and Frankenhuis, 2016). This accumulated experience is particularly important during adolescence, when key developmental processes such as synaptic pruning and myelination occur (Petanjek et al., 2011; Yakovlev et al., 1967). Both rely on repeated experience to determine which neural circuits need to be strengthened and which are less essential --in the form of experience-dependent plasticity (Dow-Edwards et al., 2019; Wilbrecht et al., 2010). In particular, the need to differentiate from caregivers and develop more independence may lead to a particular sensitivity for social experience, as indicated by rodent studies showing that social isolation in early adolescence leads to reduced spine density in the frontal cortex and changes in cortical dopamine function (Novick et al., 2011; Silva-Gómez et al., 2003; Wright et al., 2008). While these neural processes may also interact with pubertal development, some may be independent – for example, one recent study found that frontal spine pruning in rodents does not depend on the presence of gonadal hormones prior to puberty (Boivin et al., 2018). Further research, especially in animals, is needed to further identify which molecular mechanisms driving adolescent brain development actually rely on the biological changes of puberty.

Importantly, our findings suggest that the process of pubertal development appears to have a unique influence on improvements in response latency. Thus, puberty may support the optimizing of performance, while other age-related developmental processes may more specifically relate to the refinement of inhibitory control and cognitive abilities in general. In this context, performance optimization refers to the ability to modulate task responses to be quicker and more effective. This is consistent with the proposal that puberty contributes to the beginning of critical period plasticity in adolescence (Larsen and Luna, 2018) because this increased brain plasticity provides a clear mechanism through which pubertal processes help the brain to specialize and optimize its cognitive performance. Of note, the aforementioned study showing that dendritic spine pruning did not rely on gonadal hormones also found that if pre-pubertal hormones were present prior to puberty, this had an impact on the morphology of dendritic spines, suggesting a similar optimization role for puberty in dendritic spine maturation (Boivin et al., 2018). This provides further evidence that in many cases, puberty may not be essential for key neural and cognitive developmental processes but rather, it may allow the brain to hone and adjust its performance in order to reach an optimal adult endpoint.

Importantly, while canonical brain regions associated with inhibitory control were tested in this study, only VLPFC activation and connectivity were related to puberty, supporting the pubertally-dependent maturation of VLPFC as contributing to changes in latency-related aspects of behavior. Moreover, initial results on an overlapping cohort from our group looking at effects of frontostriatal connectivity (Ojha et al., this volume) also find that pubertal development is related to antisaccade latency. Since much of the literature directly linking pubertal mechanisms to regional brain development has utilized rodents, who do not have a lateral prefrontal cortex, there is a dearth of mechanistic support for how puberty might be directly implicating VLPFC. However, some human studies have shown that sex differences in the structure of the lateral PFC, such as cortical thickness, emerge at the onset of puberty (Nelson and Guyer, 2011; Raznahan et al., 2010), suggesting that puberty may have a unique influence in this area of the brain. Furthermore, VLPFC is related to stopping motor responses and response uncertainty (Levy and Wagner, 2011) as well as flexibly supporting specific inhibitory control demands depending on the task (Ryman et al., 2019). Thus, VLPFC connectivity to motor and performance monitoring (dACC) regions may support the ability to plan and execute a timely response. Accordingly, the VLPFC may be the region of the prefrontal cortex allowing for the performance optimization that we are associating with pubertal development. Activity in the cingulate region identified here, which is also known as the “mid-cingulate zone” or the “rostral cingulate motor zone” in prior literature, has previously been associated with variation in response speed (Hahn et al., 2007) and time to plan a cognitive response (Domic-Siede et al., 2021), providing a link to its association with antisaccade response latency in the current study. This area of the brain may also underlie important areas of general cognitive function such as between-network integration (Margulies and Uddin, 2019; Tang et al., 2019) and goal-directed behavior (Touroutoglou et al., 2020, 2019). Our findings, therefore, may support a potential role of VLPFC-cingulate circuitry in this puberty-related optimization of cognitive performance as contributing to pubertally-dependent changes in cognitive response speed.

Some limitations of this study should be noted. First, the age range during which puberty occurs varies dramatically across individuals based on many characteristics. Because this sample was not originally recruited with a focus on full pubertal development, resulting in relatively fewer young children, the onset of puberty may not be well-characterized in this sample, with more timepoints representing the latter half of pubertal development. Thus, we may not have been able to detect age-by-puberty interactions specific to early puberty, which are possible since the onset of puberty (transition from Stage 1 to Stage 2) represents a significant biological change. In addition, while this study took several steps to try to separate the effects of age and puberty, the fact remains that these variables still represent strongly related processes, and therefore it is still highly likely that these age and puberty effects are capturing some overlapping change. Another limitation that is common across many puberty-related studies is the fact that existing pubertal assessments reflect a number of different factors, rather than a singular measure of pubertal stage. Self-report measures, such as those used in this study, likely reflect pubertal development, but may also be affected by the individual’s self-image, their peer group, and many other factors that impact how adolescents report the status of their own development. The other commonly used method of defining pubertal status is clinician-rated Tanner stage, but this involves a physical exam that can be uncomfortable for study participants, especially adolescents. Further, puberty itself encompasses multiple neurobiological processes across both the adrenal and gonadal axes (Blakemore et al., 2010; Ladouceur et al., 2019; Marceau et al., 2015; Saxbe et al., 2015), and isolating the underlying mechanisms may require biological assays, for example by measuring hormone levels, which were not collected in this sample. Future studies should investigate whether these puberty-specific findings are associated with variation in one or multiple pubertal hormones, such as estradiol or testosterone.

## Conclusions

These findings have important implications for our understanding of puberty and its critical importance in adolescent brain and cognitive development. By elucidating the role of pubertal development in the brain, we may gain new insights into the reasons that many psychiatric disorders, and the sex differences in their prevalence, emerge in adolescence. Furthermore, this research may provide novel targets for psychiatric intervention during the adolescence years, allowing for better preventative mental health care and greater ability to treat adolescents earlier and more effectively for enhanced long-term outcomes.

